# Disentangling value, arousal and valence systems in approach-avoidance behaviors in humans using functional magnetic resonance imaging

**DOI:** 10.1101/2025.02.19.639143

**Authors:** D V P S Murty, Luiz Pessoa

## Abstract

Appetitive and aversive stimuli evoke approach and avoidance behaviors essential for survival and well-being. While affective processing has been extensively examined in terms of arousal and valence, the extent to which value processing is independent from arousal and valence processing in naturalistic contexts remains unclear. We addressed this gap using a naturalistic approach-avoidance task. Ninety-one human participants underwent functional MRI scanning while engaging in approach-avoidance tasks involving two levels of threat (mild or aversive electrical stimulation) and reward (monetary gains). We estimated effect sizes (Cohen’s D) across subjects for increasing levels of threat, reward and arousal; for valence (negative vs positive); and for valence-arousal interactions. Effect sizes for threat and reward were strongly positively correlated across brain voxels (r = 0.82), suggesting a strong influence of a shared factor. Spatial independent component analysis decomposed these effect sizes into two independent latent factors, one that represented arousal processing and another that exhibited characteristics of value processing. Importantly, we predicted that valence-arousal interaction effects would increase with latent value effects across voxels, since both valence and arousal contribute to our overall valuation process. We indeed found this to be true. Furthermore, sizable latent value effects were observed in dorsolateral prefrontal cortex, fusiform gyrus and middle temporal gyrus, areas also involved in attention and executive control. Thus, our findings revealed a value system in the human brain that could operate independently of arousal and valence systems during naturalistic approach-avoidance behaviors, providing new insights into the neural mechanisms of affective processing.

**Significance statement:** Understanding how the human brain processes affective stimuli is crucial for unraveling the neural underpinnings of behavior. Traditional models often conflate arousal and valence with subjective value, making it challenging to disentangle their independent contributions during naturalistic behaviors. In our study, we revealed independent neural correlates of value in diverse regions previously implicated in emotional processing, attention and executive control. These findings advance our comprehension of affective processing and have implications for developing targeted interventions in neuropsychiatric conditions where affective processing is disrupted.

## Introduction

Appetitive and aversive stimuli elicit approach and avoidance behavior critical for survival and well-being. Literature suggests that such affective stimuli are processed in various brain areas along multiple affective and cognitive dimensions (Pessoa, 2010, 2022; Salzman and Fusi, 2010; Anderson and Adolphs, 2014; Kragel and LaBar, 2016) such as arousal, valence and subjective value. Arousal is the intensity of emotional responses that a stimulus elicits (ranging from low to high) and valence is the hedonic quality (negative or unpleasant and positive or pleasant). Value, on the other hand, encompasses the subjective benefit or desirability of a stimulus, an action or an experience. While affective processing has been examined with a focus on arousal and valence (Russell, 1980; Posner et al., 2005), value computations have been studied extensively in the context of decision neuroscience (Kable and Glimcher, 2007; Tom et al., 2007; Rangel et al., 2008; Glimcher and Fehr, 2014) and learning (O’Doherty, 2004; O’Doherty et al., 2017). Further, value computations have been suggested to extend beyond situations that require explicit choices (Aharon et al., 2001; Bartels and Zeki, 2004; Lebreton et al., 2009, 2015; Smith et al., 2010; Tusche et al., 2010; Scheele et al., 2013). However, the extent to which value systems exist independently of arousal system in naturalistic approach/avoidance situations is not known.

Animal studies have shown the existence of diverse populations of neurons that responded to either arousal or value, in brain areas like lateral prefrontal cortex, midbrain dopaminergic areas and ventral striatum (Kobayashi et al., 2006; Matsumoto and Hikosaka, 2009; Bromberg-Martin et al., 2010; Ray et al., 2022). Consequently, population-level Blood-Oxygen-Level Dependent (BOLD) affective responses may result from the integrated effects of value and arousal arising from such sub-populations of neurons. Thus, it is imperative to delineate value and arousal systems for an accurate understanding of affective processing in humans. A few studies have attempted such a delineation, in decision and rating tasks (Litt et al., 2011; Kahnt et al., 2014). However, no study has delineated value systems from arousal systems in naturalistic contexts like approaching a reward or avoiding a predator.

In the present study, we addressed this gap using BOLD responses of 91 human subjects to 2 levels of threat and reward (high and low), each presented in different trials, in an approach/avoidance task. In a previous analysis of a subset of 80 subjects from the same dataset (Murty et al., 2023), we identified brain areas that showed significant main and interaction effects of valence and arousal using univariate analyses. In the current set of analyses, we separated independent neural representations of arousal and value processing from the observed effects of threat and reward (high vs low levels), using spatial ICA (independent component analysis (Hyvarinen, 1999)). We predicted that the neural representation of arousal should align with brain regions that are known to activate/deactivate across a diverse range of cognitive and emotional tasks irrespective of emotional valence, as documented in earlier studies (Corbetta and Shulman, 2002; Raichle, 2015). Further, valence and salience of stimuli are two important factors that contribute to overall valuation process. Value systems are known to show opposite responses to oppositely valenced stimuli (Tom et al., 2007; Matsumoto and Hikosaka, 2009; Plassmann et al., 2010), with the response magnitude depending on the stimulus salience. Thus, we predicted that the neural representation of value should resemble valence-arousal interactions in brain responses obtained through the univariate approach.

## Methods

### Subjects

Out of a total of 96 subjects (38 females) recruited from the University of Maryland, College Park, community (Murty et al., 2023), we discarded data from 5 subjects due to excessive head motion (see below) and reported results for rest of the 91 subjects (36 females) aged 21.0 ± 2.7 years (mean ± SD; range: 18-33 years). These included 44 Whites, 22 Asians, 6 African Americans and 12 whose racial origin was unspecified/multiple. All subjects had normal or corrected-to-normal vision and reported no neurological disease or current use of psychoactive drug; provided written informed consent before participating in the study; and we paid them immediately after the experiment. Institutional Review Board of the University of Maryland, College Park, approved the study.

### Experimental paradigm, data acquisition and preprocessing

The experimental paradigm, data acquisition and preprocessing steps were the same as reported in (Murty et al., 2023) and will not be repeated here. Briefly, participants controlled a turtle icon to avoid threats or pursue rewards on a screen (“play” period). There were four trial types: high and low threat (leading to aversive or mild electrical stimulation respectively if not avoided successfully), and high and low reward (leading to 100 and 10 cents respectively if pursued successfully). The result of each trial (success or failure of approach/avoidance) was indicated to the subject by a color change of the turtle icon for 1 second. Then, after a brief presentation of a blank screen (2-6 s), there was a 1-second outcome phase where participants received (or escaped/missed) either electrical stimulation or reward amounts as the case maybe. This was followed by a variable inter-trial interval lasting 6-12 seconds before the new trial began.

The participants completed the task in a single session. Each session had about 6-8 runs. Detailed description of data exclusion criteria could be found in (Murty et al., 2023). Briefly, we excluded runs that had a framewise displacement (FWD) of 4.4 mm (2 voxel lengths) or more at any time point, and runs that had 25% of all time points with FWD of 1.1 mm. First, we rejected 5 subjects who had less than four acceptable runs based on this criterion. For the 91 remaining subjects, we excluded 26 runs out of a total of 721 runs. Thus, we were left with 695 runs for analysis (mean across 91 subjects: 7.64, SD: 0.77).

### Voxel-level subject level responses

As done in (Murty et al., 2023), we used deconvolution (Friston et al., 1995) to estimate trial-averaged responses for each subject for each condition (high threat, low threat, high reward and low reward). We used *3dDeconvolve* program of AFNI suite (Cox, 1996; Cox and Hyde, 1997) (RRID:SCR_005927) to create the design matrix for every subject and fit the data to the model using *3dREMLfit* program.

Our model had the following regressors: 1) Regressors for each condition as the regressors of interest. We aligned trials to the end of the play period for each condition and modeled responses as 13 cubic splines from -10 s to 5 s relative to the play end (1.25 s interval). 2) the outcome period, modeled by convolving a 1 s square pulse with the canonical hemodynamic responses (gamma variate peaking at 4.7 s with a full-width at half-maximum of 3.77 s by using default parameters p = 8.6 and q = 0.547). 3) Our runs were of fixed duration, but the length of each trial was variable. Hence, the remaining period after the last complete trial of the run (that could have an incomplete trial or a no-task period) was modelled as a *dmBlock* basis function with 1 as the input parameter. 4) Six head motion parameters (three translational and three rotational) and their temporal derivatives. 5) Linear and nonlinear polynomial terms (up to fourth degree).

The last two nuisance regressors accounted for motion artefacts, baseline and slow signal drifts. Moreover, we censored those time points from the data whose Euclidean norm of the derivatives of the motion parameters were more than 1.1 mm (half the voxel dimension at acquisition). We removed those trials from analysis in which any data within the -10 s to 5 s window was censored (86 out of 11078 trials from analyzable runs from 91 subjects). This minimized the contribution of larger head movements to the estimates. Mean and SD (across 91 subjects and 4 conditions) of the number of trials was 30.2 and 3.56. There were at least 16 trials per subject per condition, except for 1 subject who had only 12 trials. Results did not differ significantly when we removed this subject from analysis.

We estimated the variance inflation factors (VIFs) as the diagonal elements of the inverse of the correlation matrix of the regressors (Belsley et al., 1980). This is a computationally efficient method that eliminates the need to estimate the coefficient of determination (R^2^) for regressing each of the independent regressor of the design matrix on the remaining ones. As a crucial step to inverting the correlation matrix, we also checked that the condition number was not large (this ensured that the matrix is invertible). We found that for all the subjects, the condition numbers were between 3.5 to 8.5 for every subject signifying invertibility, and the VIFs were less than 2.1, showing that multicollinearity amongst the regressors was very low.

Finally, we used for analysis only those grey matter voxels whose mean signal value (across time) was within 95-105 (signal intensity was scaled to 100 during preprocessing) and the standard deviation was less than 25 for at least 30 subjects (thus discarding 4.01% of total voxels). This step ensured that only those voxels with a good temporal signal-to-noise ratio (ratio of mean signal to standard deviation across time) for at least 30 subjects were used for analysis (N = 154297, out of a total of 160742 voxels).

### ROI-based subject level responses

For regions of interest (ROI)-based analysis, we used the 2 mm, 500 parcellation Schaefer-Yeo atlas (Schaefer et al., 2018). This atlas used a scheme of parcellation that aimed at having homogeneous resting-state functional MRI signals across voxels in each parcel, while also maintaining certain cortical areal boundaries defined by histology. This allowed us to average time-series of functionally similar voxels, yielding time-series for 500 parcels. We used non-smoothed data for this analysis. Before averaging time-series across voxels for each parcel, we removed those voxels that were also removed from voxel-level analysis. We used these time-series for estimating responses to high and low threat and reward using the same design matrix and methods as described for voxel-level subject level analysis.

### Estimation of effect sizes and residuals

We analyzed voxel-level (or ROI level) responses to each condition (high threat, low threat, high reward and low reward) from -6.25 s to 2.5 s of play end. This ensured that the early onset-associated transient was not included for analysis and also accounted for the delay in the hemodynamic response. We estimated the responses to high and low levels of arousal by averaging the responses to threat and reward for high and low levels separately. Similarly, we estimated the responses to negative and positive valence by averaging the responses to high and low levels of threat and reward respectively. We used MATLAB analysis software (RRID:SCR_001622) for all further analyses.

#### Main effects

First, we averaged the responses during the entire analysis window. We then estimated the effect sizes for threat, reward and arousal for each voxel (or ROI), defined as the ratio of the mean to standard deviation (across 91 subjects) of the pairwise differences between responses to high and low levels. Similarly, for valence, we estimated the effect sizes as the ratio of the mean to standard deviation (across 91 subjects) of the pairwise differences between negative vs positive valence responses. For valence-arousal interactions, we estimated the effect size as the ratio of mean to standard deviation (across 91 subjects) of the following pairwise differences in responses: (Threat_High_ – Threat_Low_) – (Reward_High_ – Reward_Low_). These effect sizes corresponded to Cohen’s D obtained from paired t-tests.

#### Interaction with object’s temporal proximity (period)

For the case of threat, reward and arousal, we first subtracted the low level responses from high level responses (separately for threat and reward; and averaged across threat and reward for the case of arousal). For case of valence, we subtracted the responses to positive valence from negative valence (averaged across high and low levels). Next, we split the analysis time window into two periods: mid (−6.25 to -2.5 s of play end) and late (−1.25 to 2.5 s). We then calculated the effect size for late minus mid period for each of these conditions as the ratio of the mean to standard deviation (across 91 subjects) of the pairwise differences between responses to late and mid periods. Noticeably, this corresponded to Cohen’s D obtained from a paired t-test between the late and mid periods.

For Figure 7, we regressed out the effect sizes of arousal from effect sizes of threat and reward, yielding residuals as thus:

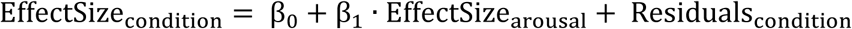

We standardized these residuals, so that these could be compared across conditions. We did not studentize the residuals as we had a very large number of voxels/ROIs. We calculated the standard deviation of the residuals as follows (Draper and Smith, 1998):

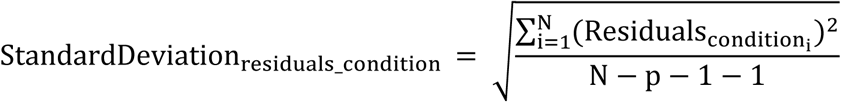

where N = number of voxels or ROIs, and p = 2 (number of regressors in the equation). We then standardized the residuals as follows:

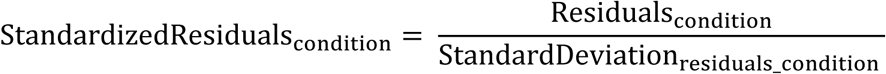

### Independent Component Analysis

Independent Component Analysis (ICA) is a mathematical method used to separate multivariate (“mixed”) signals into linearly additive and statistically independent components. In our context, ES_threat_ and ES_reward_ are observed effect sizes (Cohen’s D) for increasing levels of threat and reward respectively. According to our hypothesis, these are linear mixtures of two unknown independent factors (or components) S1 and S2. Thus, ICA decomposes ES_threat_ and ES_reward_such that

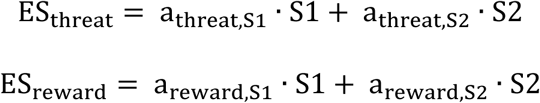

This could be represented in matrix notation as **ES_2×N_** = **A**_2×2_ · **S_2×N_**, where **ES_×N_** and **S_2×N_** are the matrices of observed signals and independent components. Since we used spatial ICA, N is the number of voxels. The matrix 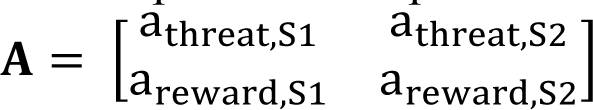 is called the mixing matrix. Its elements determine how S1 and S2 contribute to ES_threat_ and ES_reward_.

We used FastICA toolbox (Hyvarinen, 1999) in MATLAB for this decomposition. If only one of the components had a Pearson correlation coefficient |*r*|>0.5 with both *ES_threat_* and *ES_reward_*, we chose that component as the latent dominant (shared) factor (S1 in our case).

### Statistical measures

We considered Cohen’s D less than -0.5 or more than 0.5 as sizable effects (Cohen, 1988). Similarly, we considered those z-scored independent components and standardized residuals that were in the lower and upper 2.5 percentiles as sizable latent effects. We highlighted only those voxels that showed such sizable effects in the figures and text. But regardless, we used all the voxels for the analysis. We measured kurtosis for the distributions of observed and latent effects using *kurtosis.m* function. Kurtosis corresponding to normal distribution is equal to 3. We created histograms using MATLAB’s *histogram.m* function (automatic binning with uniform width across bins, but the number of bins and bin width varied across different histograms), and quantile-quantile plots using *qqplot.m* function.

## Results

### Effects of threat and reward levels did not correspond to effects of value

First, we examined the effects of increasing levels of threat and reward (high vs low levels) for each voxel in -6.25 s to 2.5 s of play end. Figure 1A shows schematic responses of arousal, valence and value systems to increasing levels of threat and reward. An arousal system would show comparable responses to similar levels of threat and reward; and its responses would increase with stimulus levels regardless of the valence. Whereas responses of valence system would be different for threat and reward but not vary across levels. Finally, responses of a value system would increase with the subjective value of the stimulus. In other words, value responses would increase from high threat to neutral to high reward. The observed BOLD effects to a stimulus would comprise of effects of arousal due to its salience combined with effects of its appetitive/aversive value. We predicted that if a voxel showed comparable effect sizes for increasing levels of threat as well as reward, the contribution from a third factor like arousal should be strong. If this was the case for many voxels in a voxel population (a small region or the entire brain), the result would be a strong positive correlation between these effect sizes across voxels (Figure 1B). Compare this prediction to the case of value systems, in which voxels showing positive effect for reward should show negative effect for threat, and vice versa (Figure 1C), implying a strong negative correlation across voxels.

**Figure 1.**
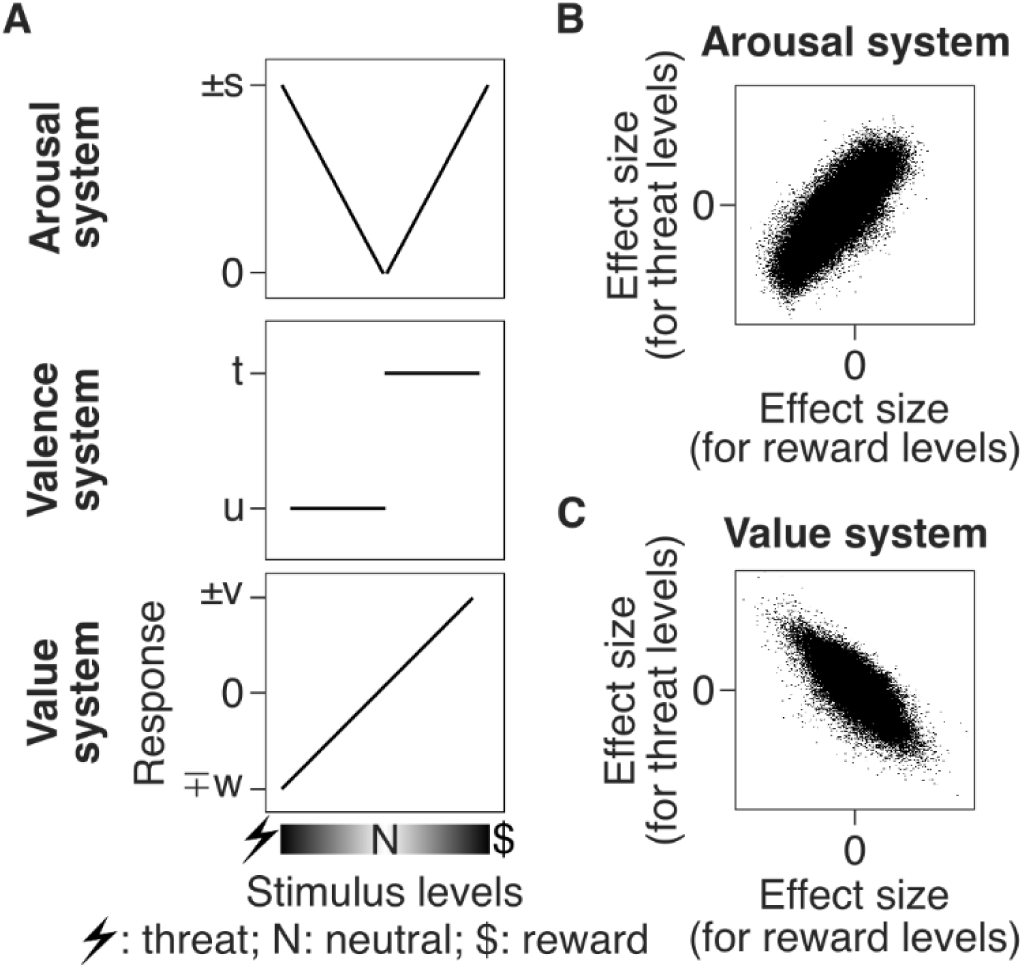
Hypotheses and goal (schematics). A) Schematic responses of arousal system (top row), valence system (middle row) and value system (bottom row) to different stimulus levels. B-C) Schematic scatter plots showing hypothesized correlation of effect sizes for increasing levels of threat and reward across all voxels: in arousal system (panel B) and value system (panel C). Each dot represents a voxel. Our goal was to find a value system in the human brain that was independent of arousal processing.

We quantified the effects (high vs low levels) of threat and reward separately using Cohen’s D (Cohen, 1988). We found that many voxels showed a sizable effect (|Cohen’s D|>0.5) for threat (23.78%) as well as reward (10.14%, Figure 2A), widely distributed across the brain. Importantly, many of these areas showed an overlap across threat and reward. Example areas included voxels in mid cingulate cortex, dorsal and ventral striatum, central amygdala, dorsolateral and ventromedial prefrontal cortex, anterior dorsal insula, frontal eye fields, intraparietal sulcus, thalamus, etc. Further, we found a strong positive correlation between these effects across voxels (*r=*0.82, Figure 2B), suggesting a strong contribution of effects from a third factor possibly related to arousal processing.

**Figure 2.**
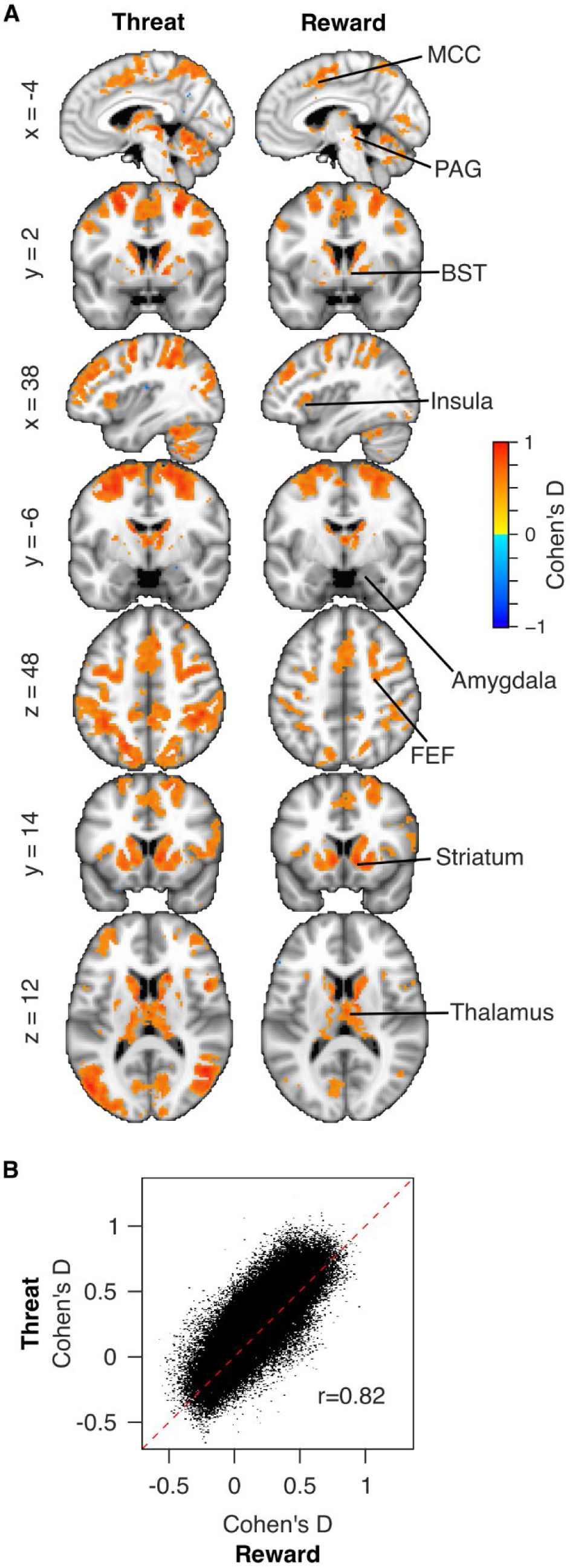
Positive correlation of effect sizes of threat and reward with each other across voxels. A) Activation maps showing sizable effects (|Cohen’s D|>0.5) for increasing levels of threat and reward. B) Scatter plot showing effect sizes for increasing levels of threat (y-axis) and reward (x-axis) across voxels (N=154297). Each dot represents a voxel. Red dashed line is the identity line. Pearson’s correlation coefficient (*r*) is indicated in the plot. BST: bed nucleus of the stria terminalis, FEF: frontal eye field, MCC: mid cingulate cortex, PAG: periaqueductal gray.

To ensure that this observation was not driven by correlations in time-series data across large sets of voxels (Fox and Raichle, 2007), we also performed an ROI-based analysis. Regions of interest (ROIs) were predefined 500 parcels in the atlas by Schaefer et al. (2018) (see Methods). We found similar results: effect sizes of threat and reward levels were highly correlated with each other across ROIs (*r*=0.87, Supplementary Figure 1).

### Decomposing the effects of threat and reward into factors representing dominant and residual effects

The above observations suggested that the observed effects of increasing levels of threat and reward were composed of two independent unknown components (or factors) S1 and S2, representing either dominant (shared) or residual latent effects. We used independent component analysis (ICA, see Methods) to decompose these observed effect sizes for all voxels into S1 and S2. It is important to note that the distribution of effect sizes of threat and reward across voxels was sub-gaussian, as revealed by their kurtosis (2.40 and 2.35) and quantile-quantile (Q-Q) plots (Figure 3A). This suggested that these represented neural processing (focused activations) and not merely noise distributed across voxels. This further suggested a priori that at least one of their components (S1 or S2) followed non-Gaussian distribution across voxels, an important requirement for ICA. This prediction was found to be true for both S1 and S2, as reflected by their kurtosis (2.27 and 3.49) and Q-Q plots (Figure 3B). Importantly, the sub-Gaussian nature of S1 suggested that this component also represented neural processing and not merely noise.

**Figure 3.**
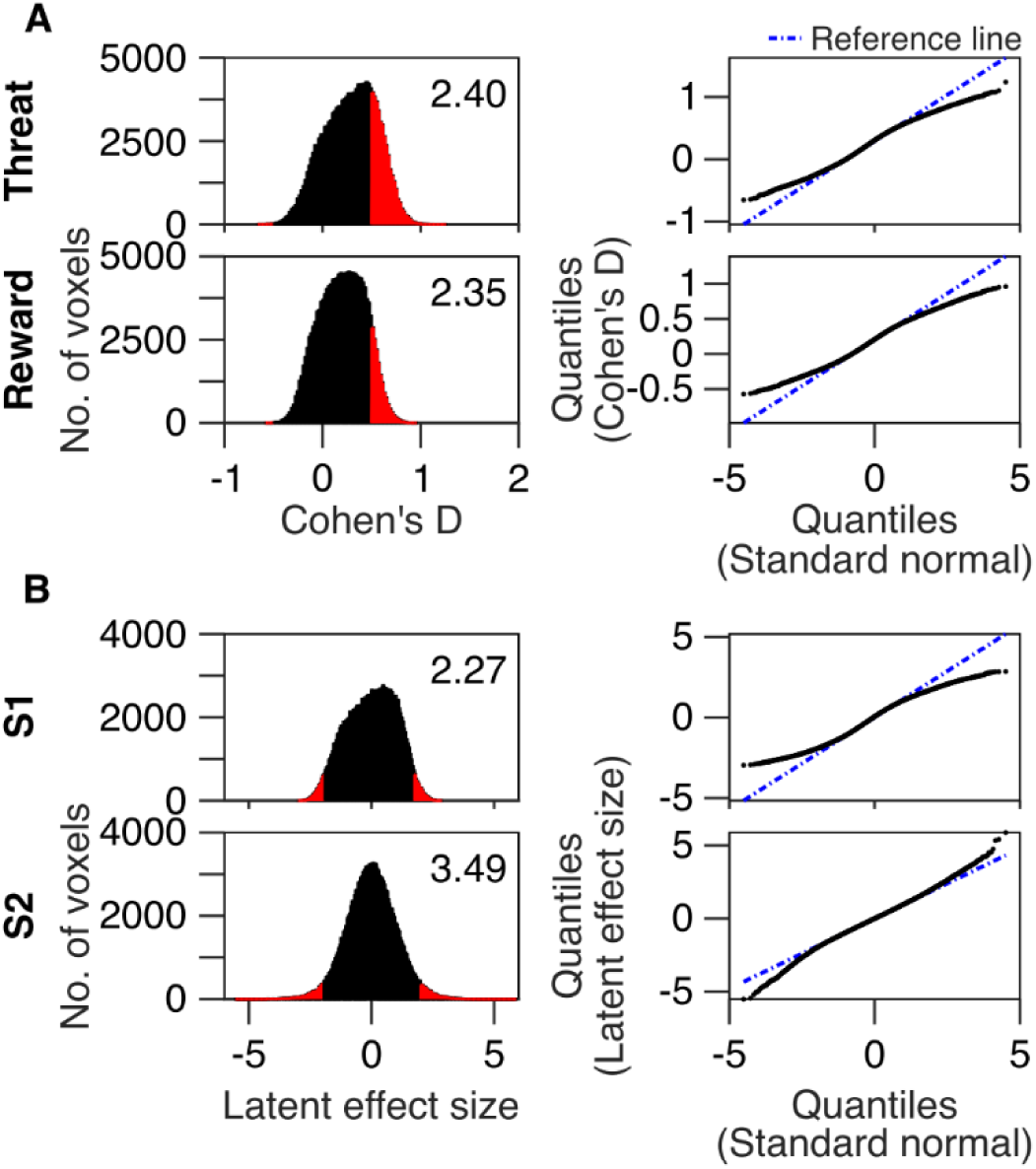
Non-gaussianity of effect sizes and independent components. Left column: histograms showing distributions of effect sizes (Cohen’s D) for increasing levels of threat and reward in panel A; and distributions of latent effect sizes for components S1 (dominant component) and S2 (residual component) in panel B. Bins representing sizable effects are shown in red (|Cohen’s D|>0.5 for threat and reward; S1<-1.91 or >1.73 and S2<-1.97 or >1.99 representing lower and upper 2.5 percentiles). Kurtosis of the distributions are indicated in the respective plots. Kurtosis for a normal distribution is equal to 3. Right column: quantile-quantile plots of the distributions in the left column. Deviation of the observed quantiles (black dots) from the reference line (blue dashed line, representing theoretical gaussian distribution) revealed the non-gaussianity of the distributions. S1 and S2 have been z-scored in this and all subsequent figures.

S1 was highly correlated with effect sizes of both threat and reward levels across voxels, explaining about 88.43% and 93.47% of variance respectively. Thus, S1 represented the dominant factor. The residual factor S2 explained the remaining 11.57% and 6.53% of variance in effect sizes of threat and reward levels across voxels. We z-scored both the components for further analysis since ICA inherently cannot determine the true scale of the decomposed components, rendering their absolute magnitude uninterpretable.

### Latent dominant factor represented arousal processing

We first determined if S1 represented arousal processing. We estimated effect sizes of arousal (Cohen’s D for high vs low levels) directly from threat and reward responses (see Methods). Using this univariate approach, we found that 32.6% of voxels showed sizable effects (|Cohen’s D|>0.5, see Figure 4A, Supplementary Figure 2 for Q-Q plot). Importantly, we found that S1 was highly correlated with the effect size of arousal, explaining about 98.6% of its variance (Figure 4A). Thus, this component captured arousal processes as predicted.

**Figure 4.**
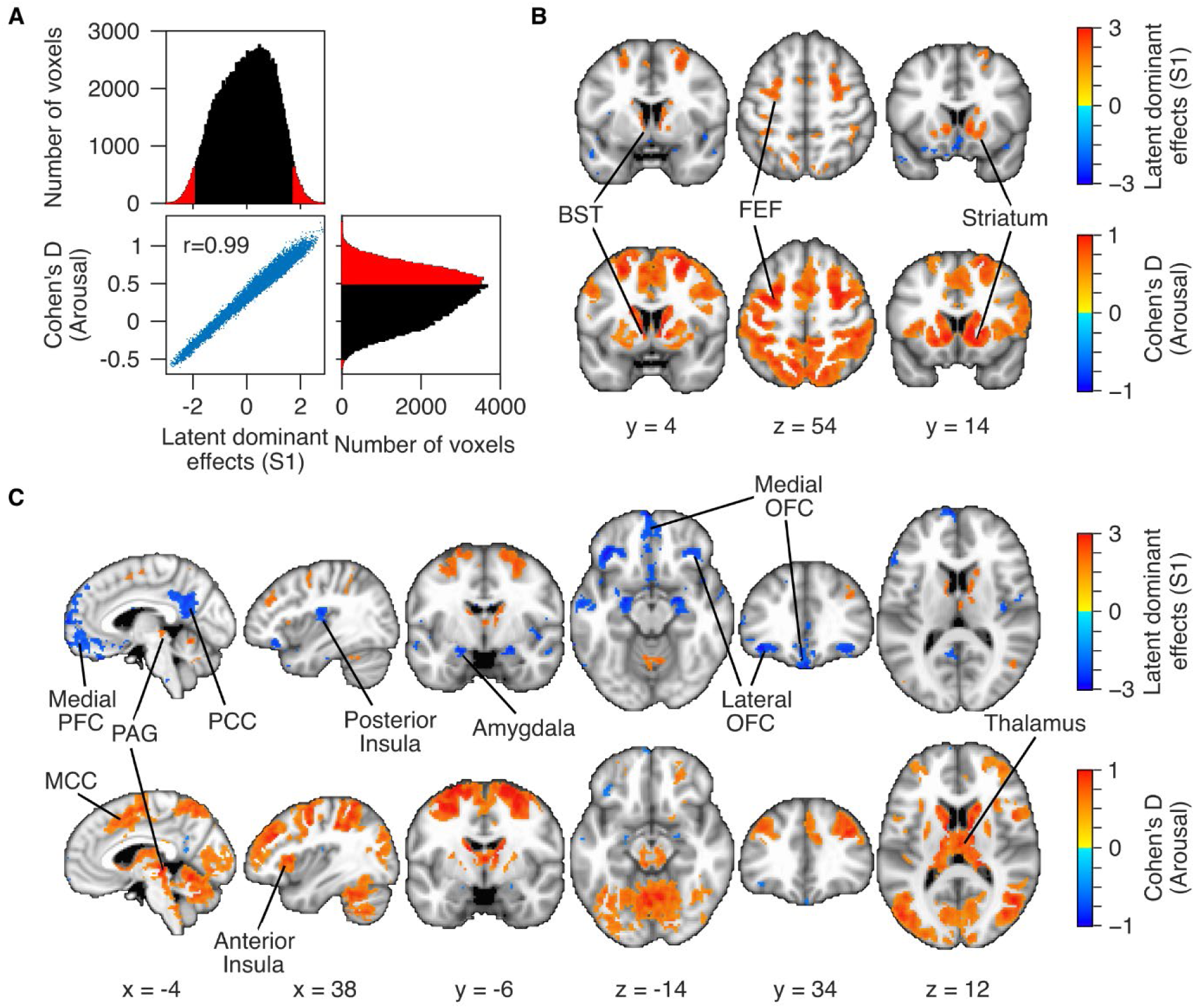
Latent dominant component (S1) and its comparison with effect sizes of arousal. A) Scatter plot showing latent dominant effects (S1, x-axis) and effect sizes for increasing levels of arousal (Cohen’s D, y-axis) across voxels (bottom row, left column). Pearson’s correlation coefficient (*r*) is indicated in the plot. Histograms show distribution of S1 (top row) and Cohen’s D for arousal (bottom row, right column) in the same format as panel A in Figure 3. B-C) Activation maps showing sizable effects for S1 (<-1.91 or >1.73, top row) and arousal (|Cohen’s D|>0.5, bottom row). Panels B and C highlight similarities and dissimilarities between the two measures. Key areas are indicated in the plots. BST: bed nucleus of the stria terminalis, FEF: frontal eye field, MCC: mid cingulate cortex, PAG: periaqueductal gray, PFC: prefrontal cortex, PCC: posterior cingulate cortex, OFC: orbitofrontal cortex.

Interestingly however, its brain representation differed from that of effect size of arousal. We plotted its z-scored values for each voxel (in the lower and upper 2.5 percentiles) on brain sections seen in Figures 4B and 4C, representing sizable latent dominant effects compared to the rest of the voxels. Areas like frontal eye fields that are known to show increases in activity to attentional processing showed positive S1 values, as well as large effect sizes for arousal (Figure 4B). Further, areas like medial PFC and PCC that are known to show task-related decreases showed negative S1 values (Figure 4C). Notably, negative S1 values were also observed in centromedial amygdala bilaterally. However, these areas were not observed to show sizable effects to arousal in the univariate maps. Further, areas such as MCC, anterior insula, etc. which showed significant effects of arousal were not captured in the brain representation of S1. In fact, we found that the conjunction (union) map of effects of threat and reward was closer to arousal map obtained through univariate approach (Jaccard coefficient: 0.72) compared to S1 (0.36), whereas S1 revealed task-positive and task-negative regions consistent with previous literature (see Discussion).

### Residual factor represented value processing

We next examined whether S2 predominantly represented value processing and not merely supra-Gaussian noise. Specifically, we tested if S2 revealed bidirectional (opposite) latent effects to increasing levels of threat and reward, characteristic of value processing. If that were the case, the effect sizes of increasing levels of threat and reward should show coefficients with opposite signs when projected onto S2 (*a_threat_,S_2_* and *a_reward,S2_* in the mixing matrix **A**, see Methods). In other words, the ratio of these coefficients should be negative. We indeed found this to be true (*a_threat,S2_*>0 and *a_reward,S2_*<0). We plotted this component’s z-scored elements on brain sections seen in Figure 5. Again, only those voxels with values in the lower or upper 2.5 percentiles have been shown in these figures, representing strong residual effects compared to other voxels. These voxels were in the striatum, dorsolateral prefrontal cortex (dlPFC), middle temporal and occipital gyri and fusiform gyrus.

**Figure 5.**
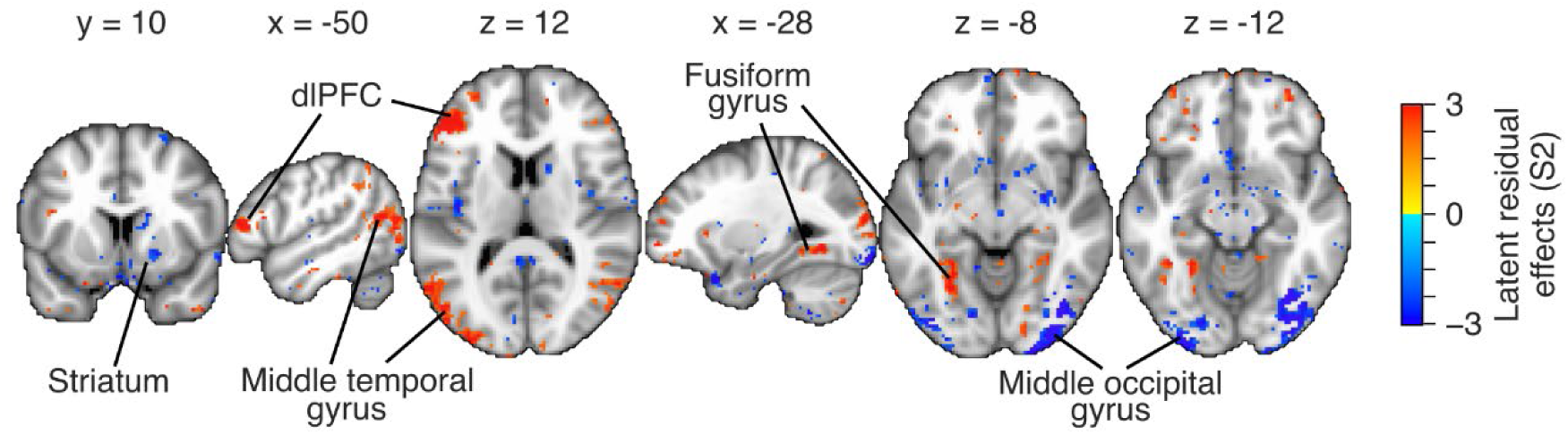
Residual component (S2). Activation maps showing sizable latent residual effects (S2<-1.97 or >1.99). dlPFC: dorsolateral prefrontal cortex.

To ensure that these results were not obtained by chance, we created bootstrapped distributions of effect sizes of threat and reward levels over 10,000 iterations. In each iteration, we chose 91 subjects randomly selected from the original set with replacement. For all these iterations, we found that effect sizes of threat and reward levels were highly positively correlated with each other (*r*=0.72 ± 0.04 (mean ± SD across iterations)). Applying ICA on these yielded dominant and residual components that were highly correlated with the original S1 and S2 reported above (|*r*|=0.95 ± 0.01 for dominant vs S1; |*r*|=0.75 ± 0.03 for residual vs S2 (mean ± SD)). Finally, the ratio of the coefficients of the residual component was negative in 99.84% iterations, consistently suggesting that this component represented value processing and not merely noise. We also ran similar tests (10,000 iterations) on simulated data. At each iteration, we simulated two vectors A and B from a Student’s *t* distribution (for non-gaussianity) such that their length and correlation matched the effect sizes of increasing levels of threat and reward. On running ICA on these vectors, we found that the ratio of the coefficients of the isolated residual component was negative in only 51% of iterations, representing noise.

### Value representation was correlated to valence-arousal interactions

Since value is itself constructed from valence and salience (arousal) of the stimulus, we hypothesized that if S2 represented value processing, it should show a strong correlation with valence-arousal interaction effects across voxels.

Operationally, we defined emotional valence of an object as negative or positive (see Methods), irrespective of its level (high or low). Many voxels in the brain (21.02%) showed a sizable effect (|Cohen’s D|>0.5) for valence (Supplementary Figure 3). These voxels belonged to anterior dorsal insula, MCC, inferior temporal and frontal gyri, lateral orbitofrontal cortex, striatum, etc. S2 explained only 1.29% of variance in effect size of valence (Figure 6A), suggesting that the neural correlates of value effects were different from valence effects.

**Figure 6.**
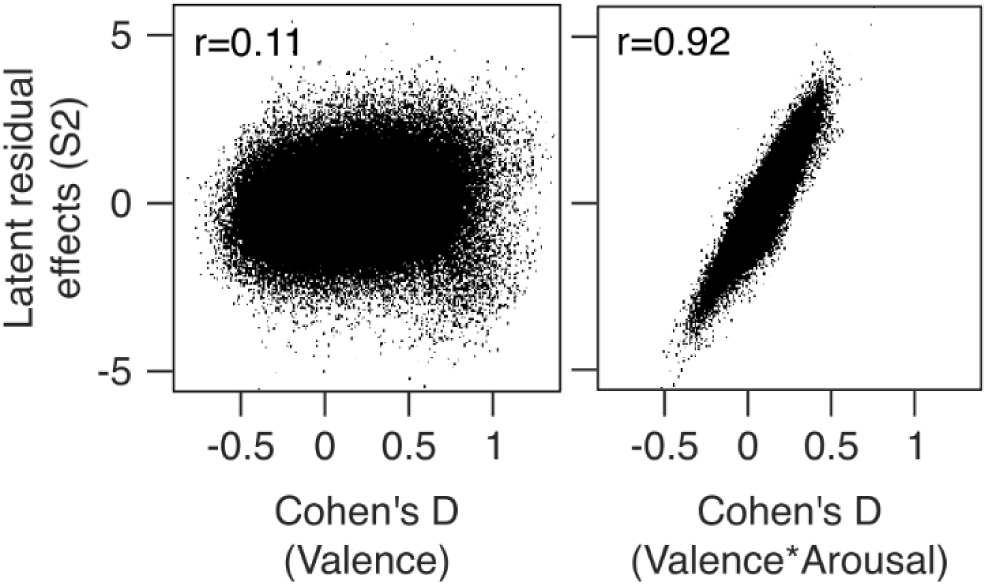
Correlation of residual component with valence effects and valence-arousal interactions. Scatter plots showing effect sizes of valence (left plot) and valence-arousal interaction (right plot) plotted on x-axis, against latent residual effects (S2) plotted on y-axis. Each dot represents a voxel (N=154297). Pearson’s correlation coefficient (*r*) is indicated in the plot.

We then estimated the effect sizes for interactions between valence and arousal (see Methods). These effect sizes were small (|Cohen’s D|<0.5) in most voxels (0.04% of voxels had sizable effects), but their distribution across voxels was non-gaussian as reflected by the kurtosis (3.37) and Q-Q plots (Supplementary Figure 4). We performed bootstrapping with 91 subjects randomly selected with replacement over 10,000 iterations. For all iterations, the effect sizes of valence-arousal interaction were highly positively correlated across voxels with the effect sizes estimated from the original set of subjects (*r*=0.76 ± 0.04 (mean ± SD across iterations)). This consistency across iterations ensured that these effect sizes, though small in magnitude, did not merely represent supra-gaussian noise.

Regardless of the size of the interaction effects, we found that S2 was highly correlated with the effect sizes of valence-arousal interactions across voxels (*r*=0.92, see Figure 6B), further supporting our interpretation that S2 represented value processing.

### Regression analysis gave comparable results for value processing

We now sought to isolate the unique effects of value through a different approach. We tested if directly regressing effect sizes of threat and reward onto the effect sizes of arousal, across voxels, yielded residuals that represented value processing (see Supplementary Information, Appendix 2 for theoretical considerations). Importantly, this approach yielded two distinct sets of residuals, one for each regression, that could potentially represent aversive and appetitive value processing separately. This was an added advantage over the ICA approach.

We plotted voxel-level z-scored residuals (in the lower and upper 2.5 percentiles) on brain sections shown in Figure 7A. As predicted, z-scored S2 was highly correlated with z-scored threat/reward residuals (variance explained by S2: 89.55% for threat and 91.27% for reward residuals). Furthermore, these residuals were negatively correlated with each other across voxels (*r*=-0.82, Figure 7B), strengthening the idea that these residuals for threat and reward predominantly represented aversive and appetitive value processing.

**Figure 7.**
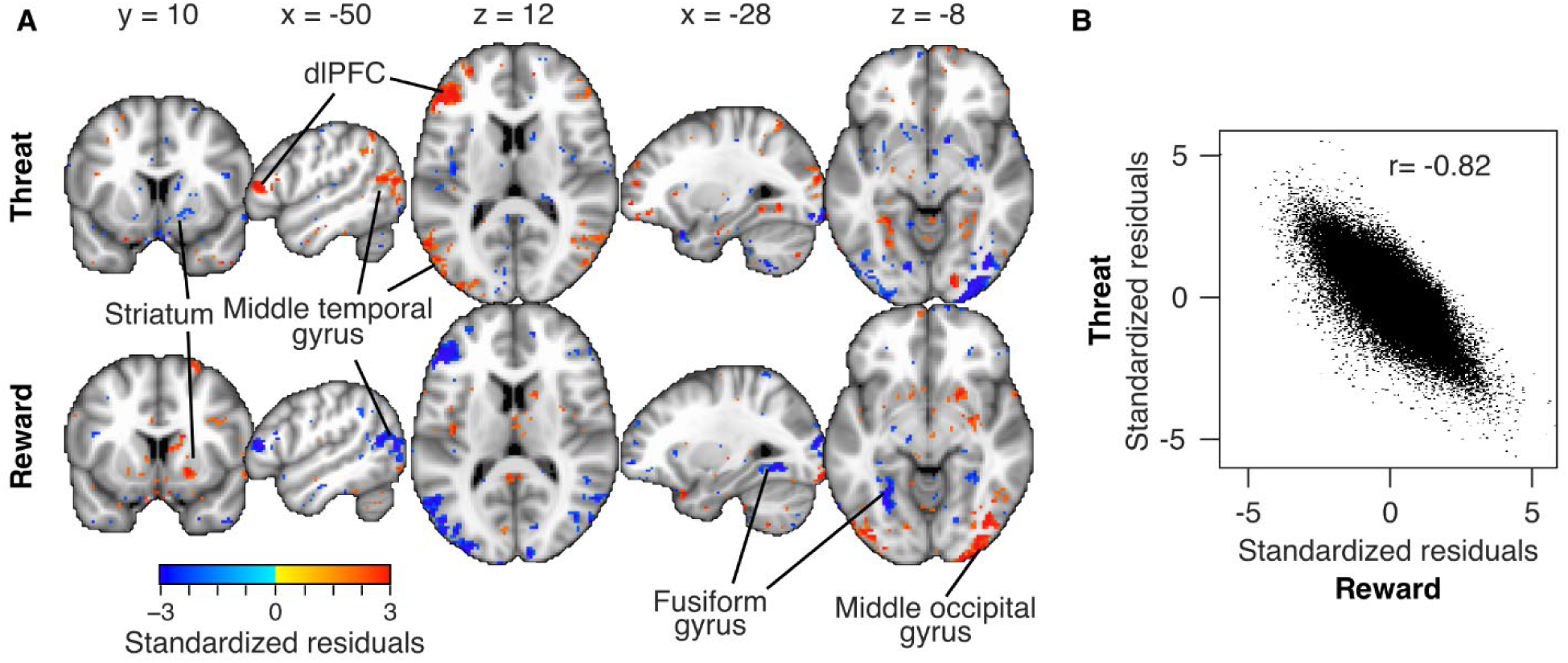
Negative correlation of standardized residuals of effect sizes of threat and reward across voxels, resembling value processing. Residuals have been obtained after regressing out the effect sizes of arousal across voxels. A) Voxels with residuals falling in the lower and upper 2.5 percentiles are shown in these brain maps. B) Residuals across all voxels are plotted as a scatter plot. Same format as in Figure 2B. dlPFC: dorsolateral prefrontal cortex.

### Temporal proximity of object had a weak effect on the value system

In our study, subjects were successful in avoiding the predator or approaching the reward in two-thirds of the trials (mean percentage ± SD across subjects: 67.52 ± 6.55%). Thus, it could be argued that the subjects could predict the positive outcome (success) of the trial as the object drew closer to the subject. This prediction may have influenced their affective responses and thus drove the positive correlation between the effect sizes of increasing levels of threat and reward. However, we ruled out this possibility by demonstrating a weak interaction of temporal proximity with threat, reward and arousal (and thus value); and also by replicating the results described in Figures 2 to 5 by restricting the analyses to -6.25 to -2.5 s of play end.

First, we explored whether arousal, valence or value effects interacted with object’s temporal proximity. We split the analysis time window into two periods: mid (−6.25 to -2.5 s of play end, farther in time) and late (−1.25 to 2.5 s, closer in time), and averaged the responses in these windows. We then estimated the effect of the interaction of threat, reward, arousal and valence with period (late vs mid period, see Methods). Noticeably, the effects of these conditions estimated in the original time window (−6.25 to 2.5 s of play end) could be regarded as main effects, as opposed to their interaction effects with period tested here. We found that only 0.03%, 0.03%, 0.09% and 4.46% of total voxels showed sizable interaction effects with period, for threat, reward, arousal and valence respectively (Supplementary Figure 5). Thus, the interaction of threat, reward, arousal and valence with period was weak in most voxels as opposed to their main effects.

Next, we restricted the analysis described in Figures 2 to 5 to -6.25 to -2.5 s of play end. We found that the effect sizes of increasing levels of threat and reward were highly positively correlated with each other (*r*=0.73). Applying ICA on these yielded dominant and residual components that were highly correlated with the original S1 and S2 (Figure 8). Finally, the ratio of the coefficients of the residual component was negative (*a_threat,S2_* >0 and *a_reward,S2_* <0). Thus, we could reproduce the value system using an earlier time window.

**Figure 8.**
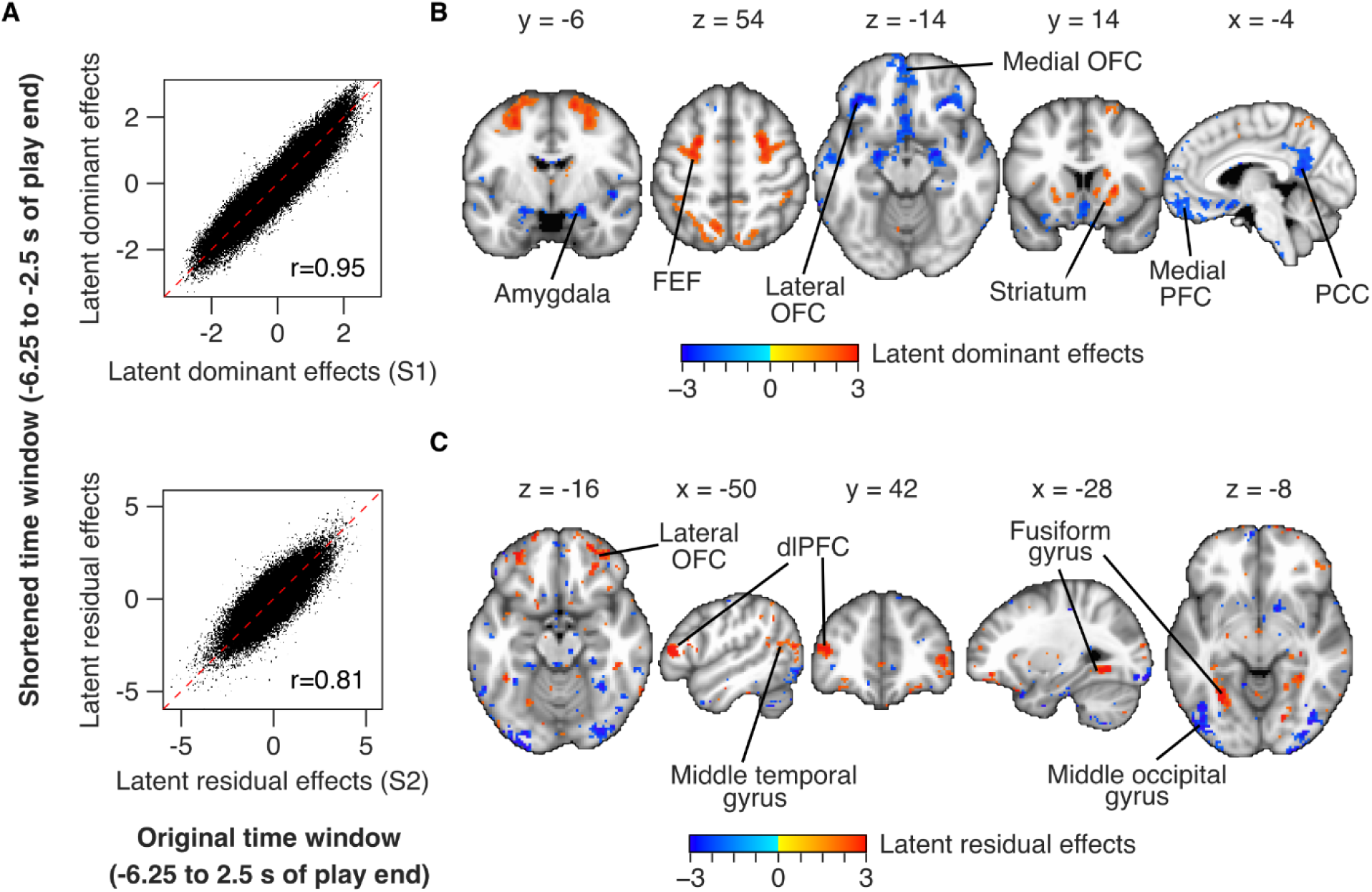
Latent effects in a shorter, earlier time-window. A) Upper plot: scatter plot showing latent dominant effects (y-axis) estimated in a shorter, earlier time-period (−6.25 to -2.5 s of play end) plotted against S1 (x-axis). Each dot represents a voxel (N=154297). Red dashed line is the identity line. Pearson’s correlation coefficient (*r*) is indicated in the plot. Lower plot: scatter plot for latent residual effects vs S2, same format as in the upper plot. B and C) Activation maps showing latent dominant and residual effects estimated in the shorter, earlier time-period. Only the dominant effects less than -1.85 or more than 1.86 have been shown in panel B (corresponding to lower and upper 2.5 percentiles); similarly, the residual effects less than -1.97 or more than 1.97 have been shown in panel C. FEF: frontal eye field, PFC: prefrontal cortex, dlPFC: dorsolateral prefrontal cortex, PCC: posterior cingulate cortex, OFC: orbitofrontal cortex.

## Discussion

In this study, using approach and avoidance situations in different trials in the same paradigm, we demonstrated that the effects of increasing levels of threat and reward on the human brain shared a dominant, arousal-related component and a residual component that tracked their distinct appetitive or aversive value. To begin with, using univariate modelling, we showed a strong positive correlation between effect sizes of increasing levels of threat and reward across voxels, suggesting a shared, salience-driven arousal process. We then used spatial ICA to decompose these effect sizes into two independent components. The dominant component resembled arousal processing as predicted. The residual component reflected bidirectional nature of value processing and was highly correlated with the effect sizes of valence-arousal interactions, suggesting that this component represented value processing. Further, we obtained comparable results for value processing when we directly regressed out effect sizes of arousal from those of threat and reward across voxels. Finally, by reproducing these results in an earlier temporal window, we demonstrated that our findings reflected the subjects’ predicted value of the object upon consumption, and not merely their prediction of success during the trial.

### Comparison with previous literature

Previous studies suggested the existence of task-positive and task-negative regions in the brain that show distinct patterns of activity during cognitive tasks (Fox et al., 2005; Fox and Raichle, 2007). Task-positive regions, such as those in the dorsal attention network (such as FEF and intraparietal sulcus) and frontoparietal control network (such as dlPFC and PPC), exhibit increased activity in a variety of affective and cognitive tasks that lead to heightened arousal (Corbetta and Shulman, 2002). Conversely, task-negative regions, primarily associated with the default mode network (including medial PFC and PCC), show decreased activity during such tasks compared to resting state (Raichle et al., 2001; Raichle, 2015). Consistent with past research, our study observed positive latent dominant effects (S1) in task-positive regions and negative effects in task-negative regions. This highlights that S1 is a more appropriate representation of arousal processing, compared to arousal effects directly estimated from threat and reward responses, which closely reflected the union of threat and reward effects.

Another important finding in our study is that centromedial region of amygdala showed negative latent dominant effects bilaterally, suggesting a decrease in affect-related activity in the imminence period that was not specific to threat and reward processing. We had observed such a decrease to sustained fear of shock in an earlier study (Murty et al., 2022). Interestingly, this effect was not revealed in the univariate analysis: we did not find sizable effects in amygdala for increasing levels of threat, reward and arousal. There is a growing body of literature highlighting inconsistencies in affective processing in amygdala in humans (Chang et al., 2015; Murty et al., 2022; Levitas and James, 2024), implicating it in aversive as well as appetitive processing, and processing related to general orienting, attention and motivational salience (Bromberg-Martin et al., 2010; Fadok et al., 2018; Steinberg et al., 2020; Yang et al., 2023). Our observations are in line with this literature.

Finally, our findings revealed sizable latent effects for value in brain regions including the striatum, dlPFC, middle temporal and occipital gyri and fusiform gyrus. These results contribute to a substantial body of literature that implicates these areas in emotional processing (Miller and Cohen, 2001; Ochsner and Gross, 2005; Kanwisher and Yovel, 2006; Delgado, 2007; Vuilleumier and Pourtois, 2007; Haber and Knutson, 2010; Sabatinelli et al., 2011, 2013; Lindquist et al., 2012). Further, middle temporal and fusiform gyri have been identified in the dorsal attention network, involved in processing of visuospatial attention (Corbetta et al., 2008; Barrett and Satpute, 2013); and dlPFC has been identified in the executive control network, involved in the control of goal-directed actions (Seeley et al., 2007; Barrett and Satpute, 2013). Whether attention and control are integrated with affective value in these areas for generating approach/avoidance behaviors needs to be tested in future studies.

### Novelty in methodological approach

The positive correlation between the observed effect sizes of increasing levels of threat and reward is an important finding that resonates with earlier literature but has not been explicitly quantified and tested before. A study examining single units in macaques found a similar correlation in activity to positively and negatively valenced stimuli (see Figure 6C in (Kobayashi et al., 2006)). In human literature, many overlapping brain areas have been shown to process increasing levels of threat as well as reward (like ventral striatum, brainstem, prefrontal cortex (PFC), insula, mid-cingulate cortex (MCC), etc. (Leknes and Tracey, 2008; Bissonette et al., 2014; Hayes et al., 2014; Pessiglione and Delgado, 2015; Levitas and James, 2024)), Thus, earlier literature suggested a priori at the observed positive correlation and motivated our analysis method.

Our analyses captured the covariance of effects of threat and reward across voxels unlike univariate methods. Importantly, our method used voxels of the entire brain. It thus maximized information arising from the heterogeneity of effects across brain regions and enabled comparison of latent effects across voxels of the whole brain. Further, our choice of ICA was motivated by our goal to isolate value effects that were independent of arousal effects. ICA has been used extensively in functional MRI literature (Calhoun and Adali, 2006; Calhoun et al., 2009) for rejection of motion artifacts (Kochiyama et al., 2005; Salimi-Khorshidi et al., 2014; Pruim et al., 2015), delineation of non-overlapping temporal networks (Calhoun et al., 2009; Li et al., 2009; Du et al., 2024), etc. The dimensions of the matrix of observed signals (see Methods) consisted of time and voxels in these applications. However, we used spatial ICA to separate latent effects of threat and reward processing. Thus, our matrix of observed signals consisted of conditions and voxels as dimensions. Non-Gaussian distribution of effect sizes across voxels (possibly due to heterogeneity and/or sparseness of effects) made spatial ICA a possibility.

### Possible models for value processing

The bidirectional nature of residual effects could be explained by two possible models of value processing. One model speculates that a single value system processes both aversive and appetitive values. The prevailing consensus in the neuroeconomics literature holds that value processing brings all stimuli onto a common currency, thereby enabling comparisons between inherently different objects like monetary gains and food (Padoa-Schioppa and Assad, 2006; Chib et al., 2009; Levy and Glimcher, 2012; Bartra et al., 2013; Lopez-Persem et al., 2020). While this has been extensively tested in terms of rewards, there has been limited evidence pertaining to weighing rewards and punishments on a common scale. Tom et al. (2007) suggested that brain areas that responded to different levels of gain in a gambling task (such as the striatum and ventromedial PFC) also responded to losses but in the opposite direction, inferring that these were coded by the same mechanism, possibly responding to the subjective value of the gambles. Similar results were observed in certain other contexts (Plassmann et al., 2010).

The other model that could be speculated is of two different systems for aversive and appetitive values (Pessiglione and Delgado, 2015), that inhibited each other, leading to the observed bidirectional effects. A related psychological phenomenon has been referred to as reciprocal activation (Cacioppo and Berntson, 1994, 1999; Larsen et al., 2003). Distinction between these two models is out of scope for this study and is a topic for future research.

### Limitation and conclusion

A major limitation inherent to our naturalistic study design was that we could not explicitly measure the responses to arousal and value. Instead, we were constrained to measure threat and reward responses, and then to estimate independent latent effects of arousal and value mathematically. We did not observe sizable residual effects (S2) in many of the affect-related brain areas like insula, MCC, amygdala, BST, ventromedial PFC, etc., even though these showed prominent effect sizes for increasing levels of threat and reward. Owing to the stated limitation, our study does not rule out the possibility of value processing in these areas but only suggests that value and arousal processing may not be separable in these areas using our method (see Appendix 1 in Supplementary Information).

In conclusion, our study provided novel insights into the distributed processing of aversive and appetitive values in approach-avoidance behaviors in human brain, distinguishing them from valence and arousal processing. Our findings highlight the importance of accounting for arousal when examining aversive and appetitive value. We believe that these results would be helpful for future studies examining the integration of value with attention and executive control in goal-directed behaviors; and for exploring new brain targets for therapies like transcranial magnetic stimulation in disorders of mood and affect.

## Acknowledgments

This research was supported by the National Institute of Mental Health (R01 MH071589 and R01 MH112517).

## Data and code availability

We have not used any unpublished code that is central to the findings reported in the manuscript. We will make the data and analysis code available to the readers upon reasonable request.

## Supplementary information

## Appendix 1

ICA requires that the source signals of the observed signals should be statistically independent. However, in our case, the predicted source signals (arousal and value) could not be assumed to be completely independent as noted in the Introduction section. In this case, ICA will still attempt to find the independent components, but with important consequences as follows:
Let observed effect sizes ES_threat_ and ES_reward_ be related to ES_arousal_ and ES_value_ such that

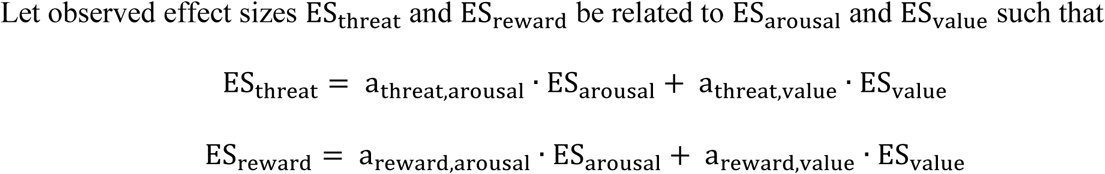

Now, since ES_value_ ∝ ES_arousal_, we could represent this as ES_value_ = β_0_ + β · ES_arousal_ + Residuals. Then the above equations could be rewritten as

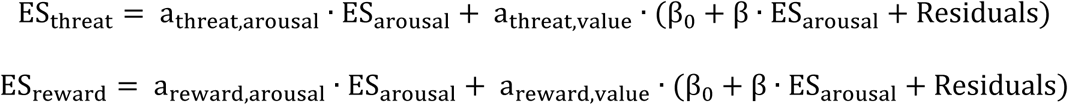

Which gives (k is a constant):

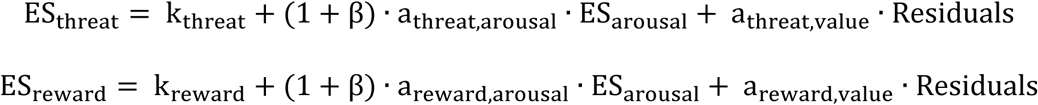

In our case, S1 was the dominant component as reflected by the variance it explained in ES_threat_and ES_reward_. Now, since ICA maximized the independence of components S1 and S2 and since ES_arousal_ and Residuals are statistically independent, we could safely assume that S1 was proportional to (1 + β) · ES_arousal_. Thus, S1 comprised of the effects of arousal as well as that portion of value which covaried with arousal and hence could not be statistically separated from arousal. But since β is a constant, S1 was simply a scaled version of ES_arousal_. Thus, z-scoring S1 reflected z-scored ES_arousal_.

However, this suggested that S2 was proportional to Residuals and not ES_value_. Thus, though S1 represented ES_arousal_, S2 represented only that portion of ES_value_that was independent of ^ES^arousal.

## Appendix 2

Given two conditions whose effect sizes are denoted by X and Y. Both these conditions have a common unknown dominant factor (with effect size) C and unknown unique factors (with effect sizes A and B), such that X = α_X,C_C + α_X,A_A and Y = α_Y,C_C + α_Y,B_B; and A and B are statistically independent from C.

There are two ways to obtain representations of A, B and C from X and Y. One way is using ICA (see Methods). The other way is to regress X and Y onto a third known variable D, such that D is highly correlated with C and acts as its proxy. Thus, D = β_D_C + ɛ_D_. Unlike ICA, this formulation is free of non-gaussianity requirements, and does not require or imply the equivalence of A and B.

In the second method, we need to test if regressing X and Y onto D yielded residuals that represent A and B. Regressing X and Y on D would yield: X = β_X_D + ɛ_X_ and Y = β_Y_D + ɛ_Y_. This gives:

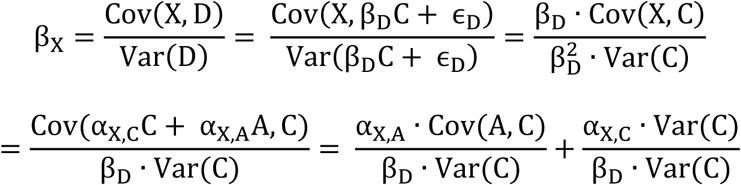

Given that A and B are statistically independent from C, Cov(A, C) = 0 and Cov(B, C) = 0

Hence, 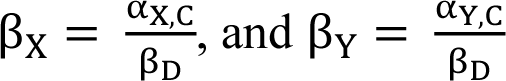

This gives: 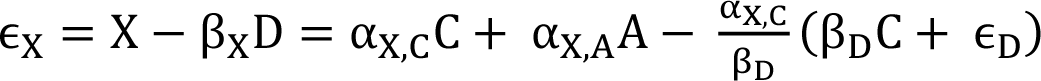

Thus, 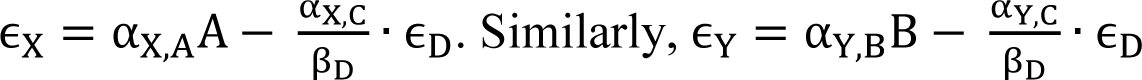

Thus, ɛ_X_ ∝ A and ɛ_Y_ ∝ B, with some added noise that reduces as the correlation between C and D increases. Thus, for residuals (ɛ_X_ and ɛ_Y_) to faithfully represent the unknown unique factors (A and B), two conditions must be met:

1. Unique factors (A and B) must be statistically independent from the dominant factor (C).
2. Correlation between the dominant factor (C) and the proxy regressor (D) should be high.

In our case, X, Y and D are effect sizes of increasing levels of threat, reward and arousal. The dominant factor C should represent arousal processing, as seen in the case of S1 component obtained from ICA analysis. Hence, C should be highly correlated with its proxy D. A and B are unique factors representing aversive and appetitive value processing (independent from arousal processing by definition) and hence statistically independent from C.

Thus, the residuals obtained by regressing effect sizes of increasing levels of threat and reward onto those of arousal faithfully represent the unique effects of aversive and appetitive value.

### Supplementary Figure legends

**Supplementary Figure 1.**
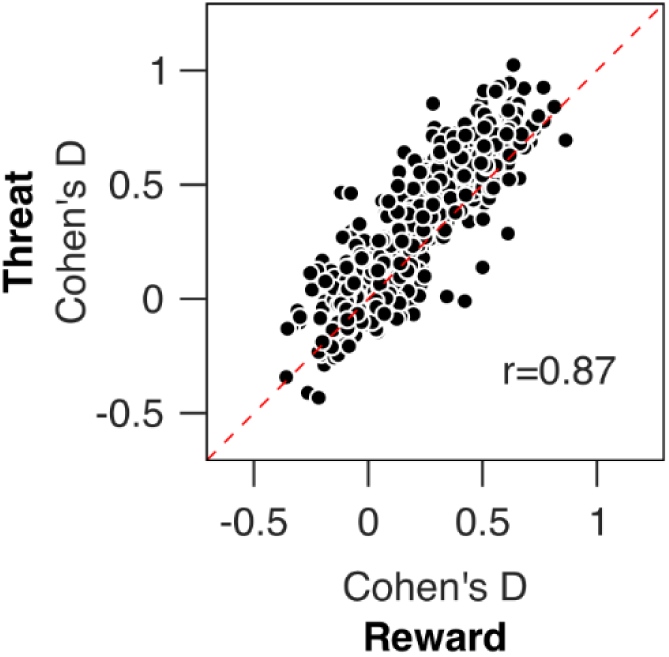
Positive correlation of effect sizes of threat and reward with each other across regions of interest. Same format as in Figure 2b, but for 500 ROIs taken from the Schaefer-Yeo atlas (Schaefer et al., 2018). Each black circle represents an ROI.

**Supplementary Figure 2.**
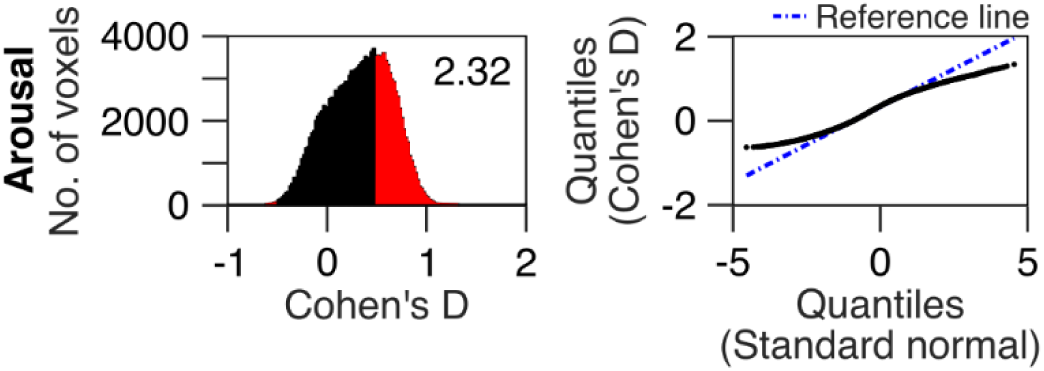
Histogram and Q-Q plots for effect size of arousal for all voxels. Same format as in Figure 3A.

**Supplementary Figure 3.**
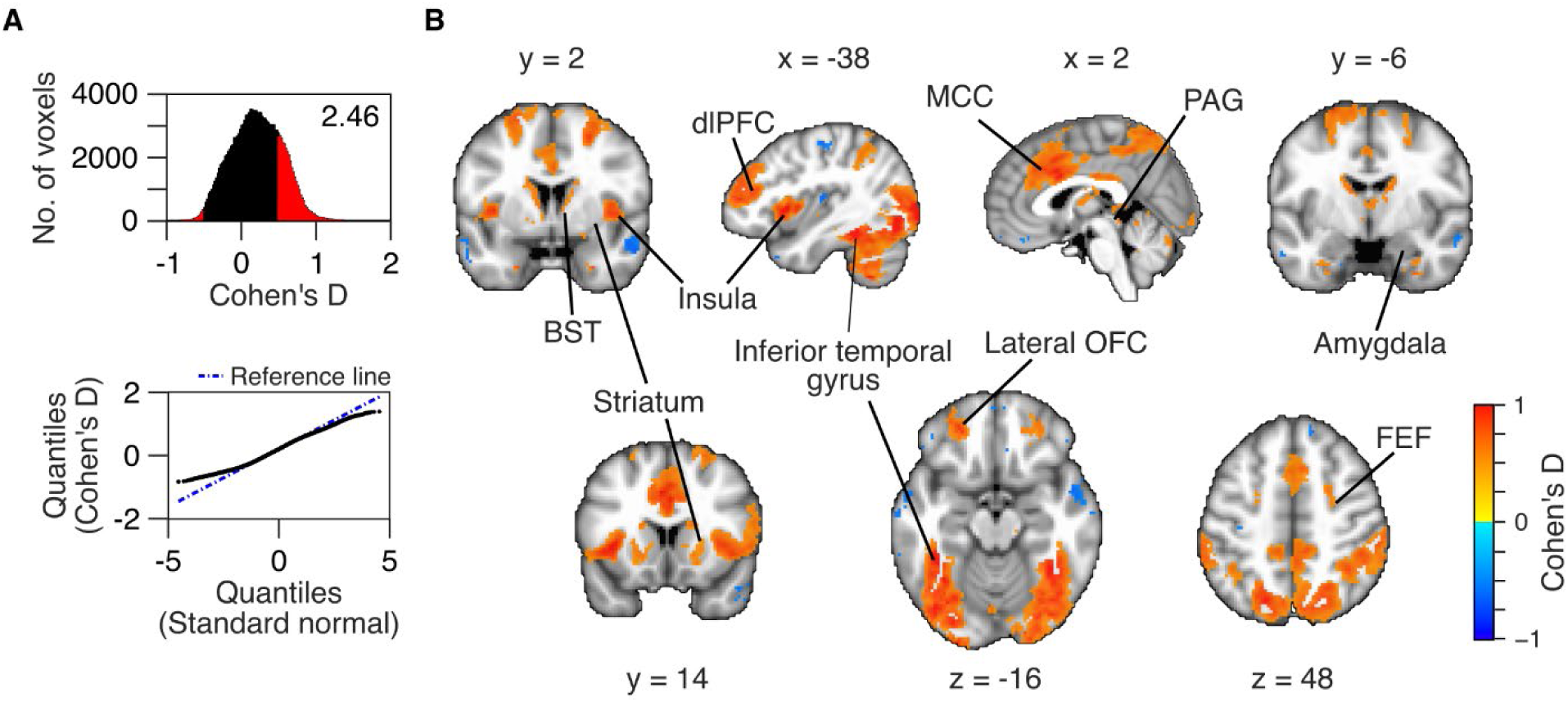
Valence effects. A) Histogram and Q-Q plots for effect size of valence for all voxels. Same format as in Figure 3A. B) Activation maps showing sizable effects (|Cohen’s D|>0.5) for valence. BST: bed nucleus of the stria terminalis, FEF: frontal eye field, MCC: mid cingulate cortex, PAG: periaqueductal gray, dlPFC: dorsolateral prefrontal cortex, OFC: orbitofrontal cortex.

**Supplementary Figure 4.**
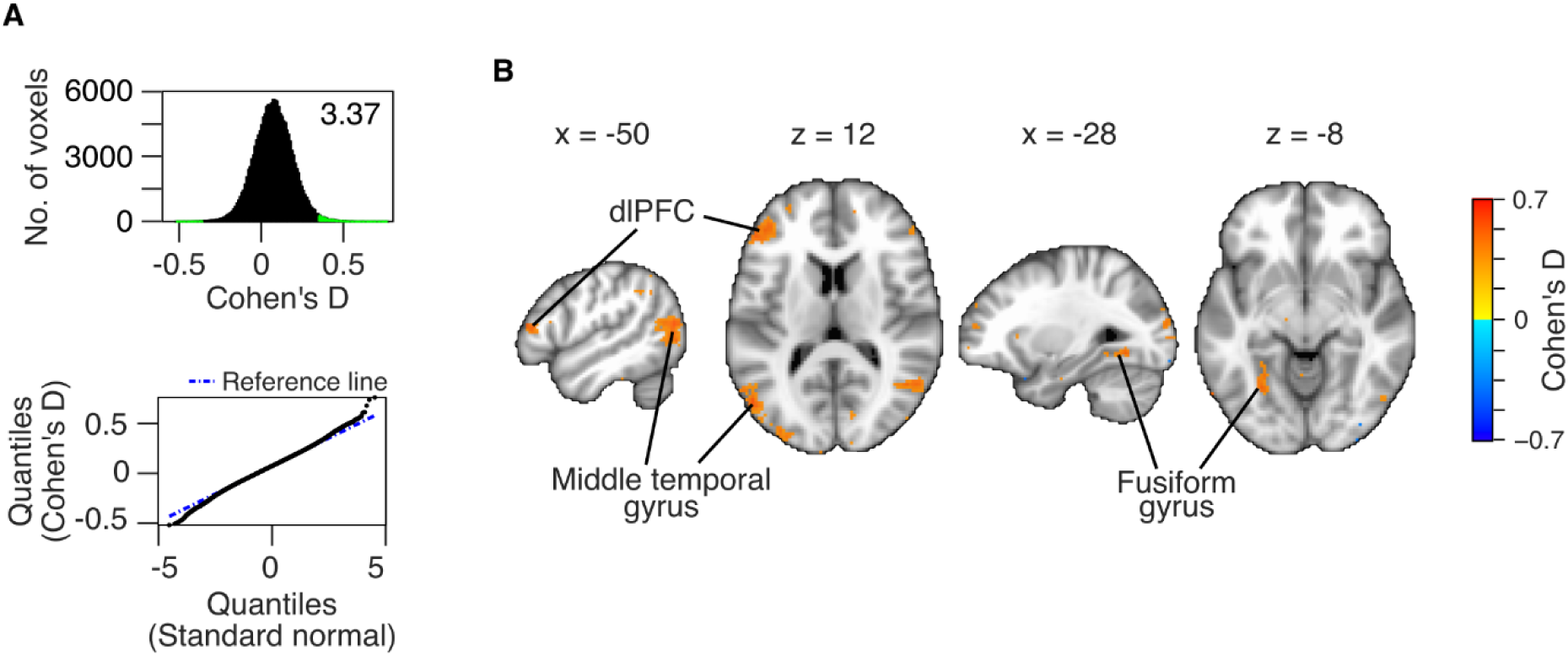
Valence*arousal interaction effects. A) Histogram and Q-Q plots for effect size of valence*arousal interaction for all voxels. Note that the effect sizes were small for a majority of voxels. Hence, we reduced the threshold (|Cohen’s D|>0.35) and marked the bins that satisfied this threshold in green. As in Figure 3A, kurtosis of this distribution is indicated in the histogram plot, and blue dashed line in the Q-Q plot is the reference line representing theoretical gaussian distribution. B) Activation maps showing |Cohen’s D|>0.35 for valence. dlPFC: dorsolateral prefrontal cortex

**Supplementary Figure 5.**
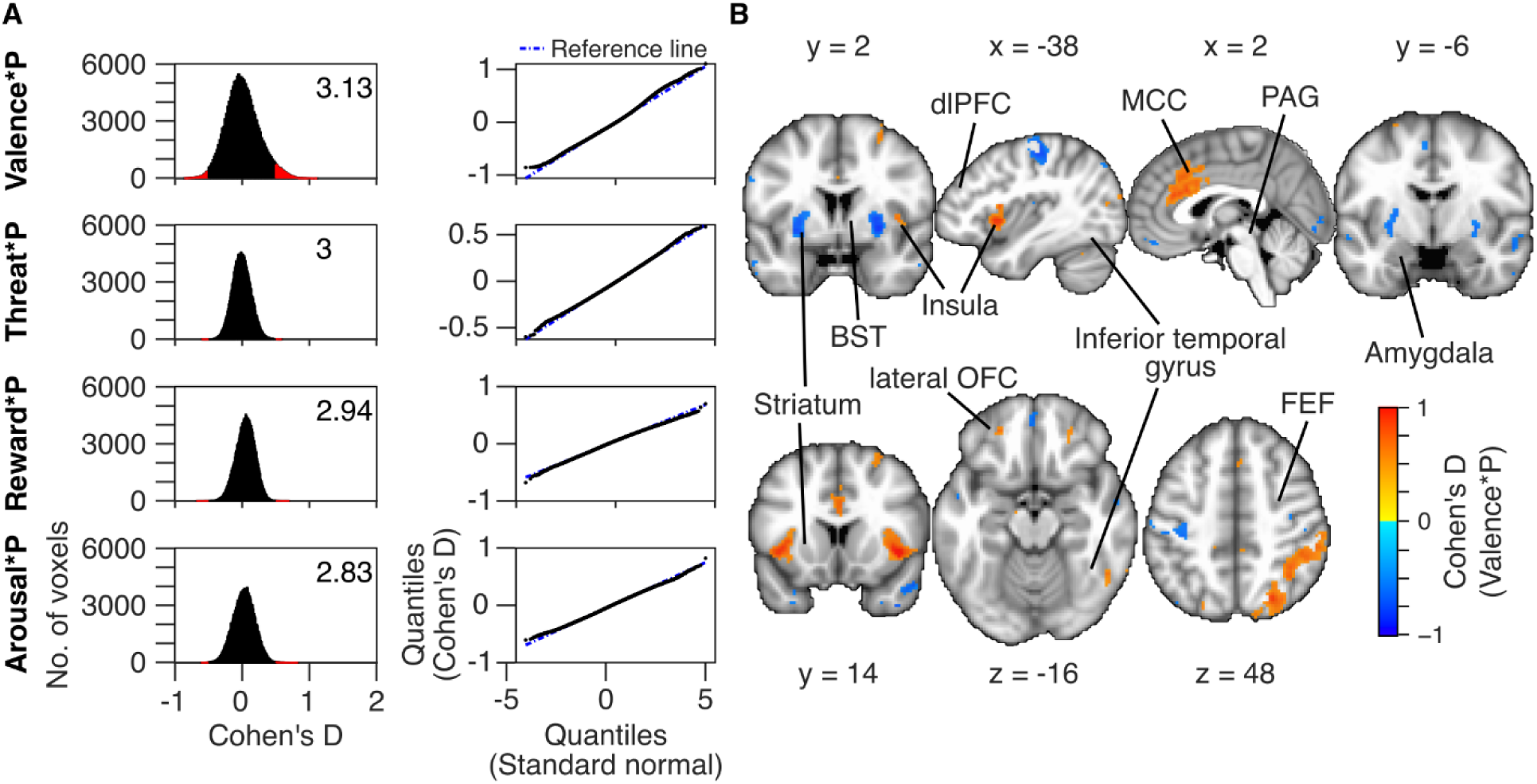
Interactions with object’s temporal proximity. A) Left column: histograms showing effect sizes (Cohen’s D) for interaction of threat/reward, arousal and valence, with period (−1.25 to 2.5 s versus -6.25 to -2.5 s of play end). Right column: corresponding Q-Q plots. Bins representing sizable effects (|Cohen’s D|>0.5) are shown in red. Kurtosis of the distributions are indicated in the respective plots. B) Same as in Supplementary Figure 3B, but for valence*period interaction. Notation ‘*P’ in the plot labels indicates interaction of a given condition (threat, reward, arousal or valence) with period. BST: bed nucleus of the stria terminalis, FEF: frontal eye field, MCC: mid cingulate cortex, PAG: periaqueductal gray, dlPFC: dorsolateral prefrontal cortex, OFC: orbitofrontal cortex.

